# Automated segmentation by deep learning of neuritic plaques and neurofibrillary tangles in brain sections of Alzheimer’s Disease Patients

**DOI:** 10.1101/2023.10.31.564976

**Authors:** Lea Ingrassia, Susana Boluda, Gabriel Jimenez, Anuradha Kar, Daniel Racoceanu, Benoît Delatour, Lev Stimmer

## Abstract

Alzheimer’s Disease (AD) is a neurodegenerative disorder with complex neuropathological features, such as phosphorylated tau (p-tau) positive neurofibrillary tangles (NFTs) and neuritic plaques (NPs). The quantitative evaluation of p-tau pathology is a key element for the diagnosis of AD and other tauopathies. Assessment of tauopathies relies on semi-quantitative analysis and does not consider lesions heterogeneity (e.g., load and density of NFTs vs NPs).

In this study, we developed a deep learning-based workflow for automated annotation and segmentation of NPs and NFTs from AT8-immunostained whole slide images (WSIs) of AD brain sections. Fifteen WSIs of frontal cortex from four biobanks with different tissue quality, staining intensity and scanning formats were used for the present study. We first applied an artificial intelligence (AI-)-driven iterative procedure to improve the generation of pathologist validated training datasets for NPs and NFTs. This procedure increased the annotation quality by more than 50%, especially for NPs when present in high density. Using this procedure, we obtained an expert validated annotation database with 5013 NPs and 5143 NFTs. As a second step, we trained two U-Net convolutional neural networks (CNNs) for accurate detection and segmentation of NPs or NFTs. The workflow achieved a high accuracy and consistency, with a mean Dice similarity coefficient of 0.81 for NPs and 0.77 for NFTs. The workflow also showed good generalization performance across different patients with different staining and tissue quality. Our study demonstrates that artificial intelligence can be used to correct and enhance annotation quality especially for complex objects, even when intermingled and present in high density, in brain tissue. Furthermore, the expert validated databases allowed to generate highly accurate models for segmenting discrete brain lesions using a commercial software. Our annotation database will be publicly available to facilitate human digital pathology applied to AD.

## Introduction

Alzheimer’s disease (AD) is histologically characterized by the occurrence of two pathognomonic brain lesions: extracellular plaques mainly composed of amyloid-ß peptides and neurofibrillary tangles (NFTs) made of abnormally phosphorylated tau aggregates within neuronal soma and dendrites/axons. The association of amyloid deposits with a peripheral crown of dystrophic tau-positive neurites forms the so-called neuritic plaque (NPs).

The standard, consensus-derived, neuropathological staging of AD relies on an ABC score [22] combining A) the topographical distribution of amyloid plaques (Thal’s phases [35]), B) the dissemination of phosphorylated tau across cortical brain areas (Braak staging, [3, 4]) and C) the neocortical density of NPs ([21]). ABC scoring relies on a binary algorithm (reaching or not specific Thal’s phases or Braak’s stages with 6 degrees of freedom for each scale) and on semi-quantitative manual assessment of NPs (CERAD criteria: from none to frequent density).

There are several questions and limitations related to standard ABC scoring, both methodological and conceptual.

First, while being instrumental in staging AD neuropathological changes, the ABC score clearly presents some limits. Hence, manual inspection and quantification of various brain regions remains time-consuming and associated with a possible suboptimal inter-rater reliability [1, 2]. It is expected that the rise of digital pathology combined with artificial intelligence (AI) algorithms could offer new solutions to allow faster and more reproducible quantifications of AD brain lesions.

Second, considering more fundamental aspects, ABC staging can hardly be used to describe and quantify fine-grained neuropathological changes such as variable NFTs and/or NPs densities [27] that may actually correspond to different pathological entities. Also, the clinico-pathological diversity of the AD spectrum [11, 14, 23] involves different variants or clusters of brain lesions with discrete anatomical localizations that may tentatively be appreciated through the study of their spatial distribution (e.g., typical AD vs limbic predominant, hippocampal sparing, behavioral/dysexecutive or posterior cortical subtypes). However, these spatial “macroscopic” descriptors may be paralleled by more subtle alterations at the “microscopic” scale that could reveal additional information to further stratify AD variants. There are indeed subtypes of the disease (e.g. rapidly progressive AD cases [31, 32]) that do not present overt topographical differences with classical AD in terms of lesion distribution. The detailed and quantitative appreciation of the different brain lesions in these cases cannot be fully appreciated through standard and manual routine neuropathological examination.

Third, the driving impact of tau lesions on neurodegeneration and on the emergence of symptoms has been repeatedly demonstrated, in both postmortem studies [24] and in vivo analysis [17] and the regional burden of tau lesions closely associates with the various AD clinical variants [27]. Tau pathological aggregates, even when analyzed in AD brains, have diverse and partially overlapping morphological phenotypes (e.g., NFTs, NPs, tortuous fibers etc.) that are not classically discriminated following standard neuropathological assessment (lack of tools to automatically count the different lesions).

The possibility to detect and segment somatic NFTs and NPs might therefore be of high importance to stratify AD individuals. Nelson and colleagues [25] hence identified a minority (10%) of “tangle-intensive” patients presenting high NFTs densities with a paucity of NPs that contrasted with “plaque-intensive” cases displaying intermediate severity NFTs but high density of NPs. More recently it was demonstrated that different tau species are present in NFTs and NPs and are associated with distinct properties in terms of seeding activity and synaptotoxic effects [30]. Interestingly, the debated “primary age-related tauopathy” (PART) entity is described as a NFT-predominant condition [12], separated from AD or, alternatively, classified as “part” of the AD continuum [6]. It is hence tempting to consider that NFTs accumulation in the medial temporal lobe of aged mild-cognitively impaired individuals, when subsequently combined with emerging NPs, triggers a quantum leap in the neurodegenerative progression.

All these data reveal an unmet need in modern AD neuropathology related to the detection-segmentation and quantification of discrete lesions (such as tau lesions that rely on multiple heterogeneous morphologies).

Recent research effort was implemented to develop automated histological classification of the different tau lesions, mainly (but not only) through AI-based approaches and virtual histological preparations (whole-slide images -WSI). Using color deconvolution and object detection algorithms Irwin and colleagues [10] were, for instance, able to automatically quantify tau cytoplasmic inclusions. More recent attempts were made to quantify by AI tangle burden using deep learning (DL) and weakly supervised learning models [34, 37], but focus was only made on tangle detection and not on their fine segmentation. Wurtz et al. [36] used fully convolutional neural networks to segment NFTs but did not provide extensive morphological analysis. The main difficulty in process automation lies in the segregation, by algorithms trained in “complex and heterogeneous” tissue environments, of the different tau-positive morphologies (e.g., discrimination between NFTs vs NPs patches). Neltner and colleagues [26] used Aperio ScanScope XT image analysis software and pattern recognition tools to quantify tau burden; the authors reported a good correlation between automatic tau burden segmentation and manual estimation of overall tau load but individual object isolation was not possible. Based on Google AutoML algorithms, Koga and Dickson [15] performed automatic detection of tau-positive tufted astrocytes, astrocytic plaques and neuritic plaques but did not investigate the potential of their models to detect and segment NFTs. The same authors succeeded in another published work [16] to classify lesions occurring in different tauopathies including cytoplasmic NFTs and coiled bodies. However, their efforts were directed to detection and classification while no information was given on the segmentation of the different objects.

It appears, therefore, that automatic detection through (deep) machine learning of the different lesion morphologies occurring in tauopathies and particularly in AD is an emerging and burgeoning field of research. Our previous work introduced novel open-source methods for quantifying NFTs and NPs based on our own studies and developments [13, 20]. These methods were recognized as pioneering contributions in recent medical image computing conferences [13], but still lead to suboptimal performances.

Therefore, in the present work, we first developed, using a widely distributed and user-friendly software (Visiopharm®), new strategies to optimize and fasten human annotations while preserving quality of analysis by setting up a “human-in-the-loop” pipeline that may reduce the annotation burden when preparing data training sets (ground truth databases) to be used in DL models.

We then used the generated database to build models for NPs and NFTs segmentation allowing to segregate, for the first time, these two categories of discrete histological objects in an automated and unbiased way.

The engineered tools may subsequently be used to accelerate, refine and make the histological diagnosis more robust, but will also be used to uncover and quantify subtle morphological differences between AD variants that otherwise could not be neuropathologically stratified.

## Materials and Methods

### Patient Selection

The brain sections used in this study were kindly provided by four different brain banks, including: Neurological Tissue Bank-Biobank-IDIBAPS, Hospital CLínic-IDIBAPS, Barcelona, Spain; IOP London Institute of Psychiatry, King’s College Hospital, Department of Clinical Neuropathology, London, UK ; Queen Square Brain Bank, Institute of Neurology - University College London, London, UK; The French National Brain Bank Neuro-CEB, Pitié-Salpêtrière Hospital, Paris, France. The use of all specimens for the present research was approved by the local ethics committee and written informed consent was obtained from all participants or their legal representatives, in accordance with the ethical standards of the 1964 Declaration of Helsinki. Paraffin embedded brain tissue sections were available from patients with sporadic AD (n=10) and patients with genetic AD (APP duplications or APP point mutations; n = 5). Cases were selected according to the following criteria: (i) pathological: well-characterized clinical cases with confirmed postmortem diagnosis of AD and minimal or absent neuropathological comorbidities (**Supplementary table 1**); (ii) technical: automated staining of tissue presenting usual intrinsic variability (in terms of postmortem preservation, fixation, coloration intensity) as observed in routine histopathological practice.

### Immunohistochemistry

Immunohistochemistry was performed on the formalin-fixed paraffin-embedded (FFPE) tissue sections (3-5μm) from frontal cortex (BA 8/9) using the Nexes station automated system (Ventana Medical System Inc., Roche). Deparaffinised and rehydrated sections were pretreated and blocked, then incubated with the AT8 primary antibody (clone AT8, monoclonal, ThermoScientific, 1:500 dilution), followed by the detection with the ultraView Universal DAB Detection kit (Ventana Medical System Inc., Roche), according to the automated procedure. The kits include secondary antibody (goat anti-mouse IgG) conjugated with horseradish peroxidase (HRP). The addition of H202 and diaminobenzidine leads to the formation of a brown precipitate in presence of HRP. Tissue sections were counterstained with hematoxylin.

### Slide digitization and annotation

All sections were digitized to obtain whole slide images (WSIs) using two scanners: Hamamatsu NanoZoomer S60 (n = 4 cases digitized) and Hamamatsu NanoZoomer 2.0-RS (n = 11 cases). Images were acquired at 200x magnification and saved in ndpi format.

All WSIs were uploaded to the Visiopharm analysis software (Visiopharm®, v2022.02, Hørsholm, Denmark) that was used for object annotation, detection-segmentation, review, and correction. Neuritic plaques (NPs) and neurofibrillary tangles (NFTs) in neuronal somata were annotated separately within regions of interest (ROI) of the cortical grey matter. All objects (NFTs or NPs) were outlined manually using Visiopharm’s integrated drawing tools. All annotated objects were verified and validated by the expert annotators (LI, LS, SB).

A neuritic plaque was operationalized as an object of a rounded to ovoid shape with a crown formed by dystrophic (enlarged, strongly AT8-positive) neurites and a central fibrillary core mostly devoid of AT8 immunoreactivity. In addition to this, we added objects as NPs with less visible or no central AT8-negative core but with clearly visible dystrophic neurites of circular arrangement. Dystrophic neurites loosely distributed in the neuropil with no specific circular arrangement were included in the “non-NPs background” category.

A tangle-bearing neuron was outlined as a neuron containing a visible nucleus and cytoplasmic fine granular, coarse granular, or fibrillary/condensed AT8 immunopositivity.

Other AT8-positive structures including neuropil threads, neuropil granules/grains, and ambiguous non-neuronal phospho-tau staining were categorized as “non-NFTs background”.

### Ground truth dataset built by iterative AI-driven object annotations

The identification of all objects of interest within a given region is often challenging for a human annotator especially within heavily stained tissue areas. Thus, following the initial human annotation of NPs and NFTs, all annotated slides were submitted to an iterative AI-driven, human-in-the-loop, object correction procedure in order to find “overlooked” or “wrongly-annotated” objects by the experts and/or to refine their outlines (**Figures 1-2)**. For this purpose, an initially annotated data set performed by human experts was used to train a U-Net convolutional neuronal network, which was a part of the AI-Architect module of Visiopharm image analysis software. The algorithm relied on 1024x1024 pixels sized training windows as well as several data augmentation steps including image rotation and flips, and hue and intensity variations.

**Figure 1:**
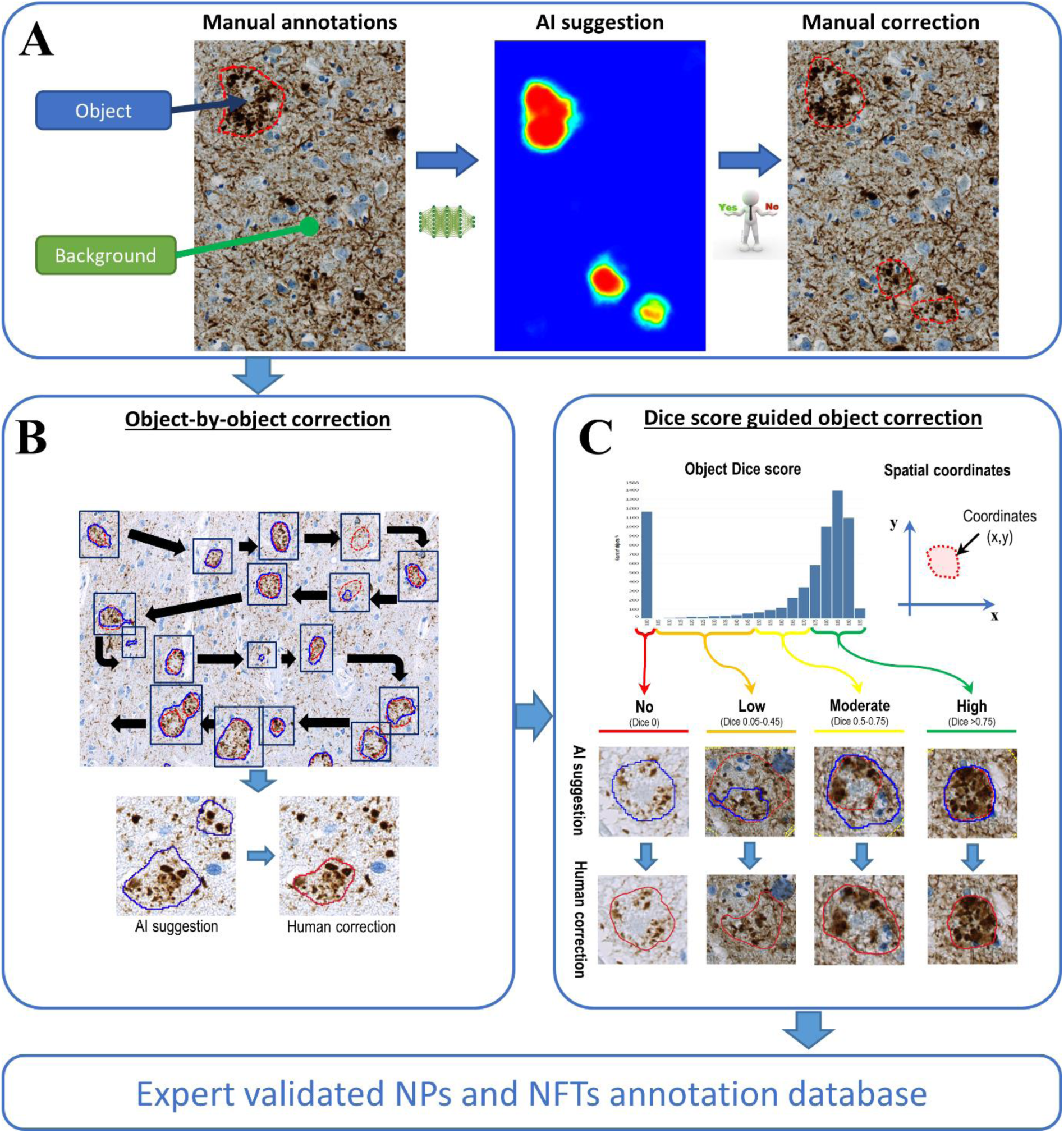
AI-driven iterative procedure for the improvement of annotation accuracy. **A:** The Human annotator outlines the object of interest (red dashed line surrounding a NP). The other part of the image is considered as “Background” by the AI algorithm. After the training, AI suggests the presence of possible non-previously annotated objects of interest (middle panel: detection probability map with red = highest probability and blue = lowest probability). The human annotator then decides for each object, if AI-outlined object is an object of interest or not and confirms/modifies the annotation boundaries if necessary (here, two red outlined small NPs were clearly overlooked by the expert and should be added to the database; bottom of the right panel). **B:** Workflow for the object-by-object improvement of the annotation dataset. Initial human annotations (red outlined objects) and AI segmentation suggestions (blue outlined objects) are delineated. A human corrector zooms in each individual object and corrects the outline if necessary and then moves to the next object. **C:** Workflow for the Dice score guided object correction. Individual Dice scores (distribution visualized as an histogram) and spatial coordinates are calculated for each individual object. Objects with a “0” or low Dice score are identified in a list of detected objects and highlighted based on their spatial coordinates. These objects can be zoomed individually (upper row: AI suggestions in blue and human annotations in red), verified, and corrected (lower row: human corrections in red). This procedure allows the selection of a subcategory of “problematic” objects (e.g., objects with “0” Dice score) for more targeted and efficient corrections.

**Figure 2:**
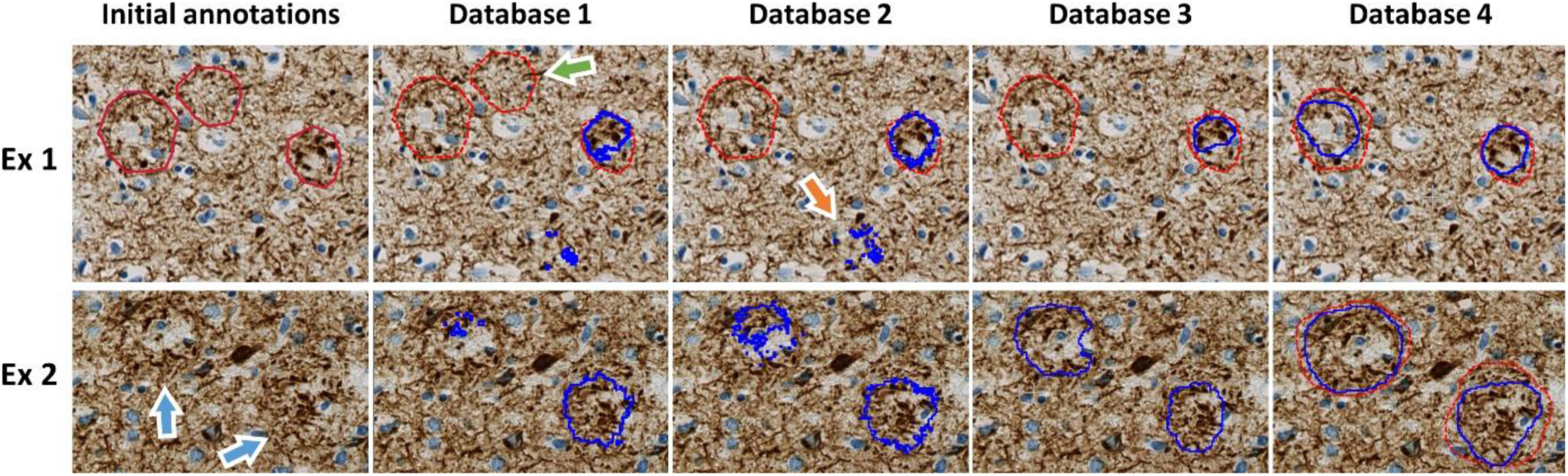
Two examples (rows Ex1 and Ex2) of gradual improvement of the annotation quality of NPs using the object-by-object correction method. Initial annotations were done by human expert (red outline in first column) and certain objects were clearly “overlooked” by experts (blue arrows in Ex2). These initial annotations were used as ground truth for AI training. The trained model generated its own set of segmentations (blue outlines in second column). During subsequent iterative AI modeling steps all annotations were reviewed by pathologists (Human-in-the-loop) and objects without sufficient agreement between experts were removed (green arrow in Ex1). The overall procedure leads to disappearance of false positive objects detected by AI (orange arrow in Ex1) and to a gradual convergence of human and machine segmentation (last column).

The initial phase of object correction involved three cycles of AI training and object-by-object validation for NPs and NFTs (**Supplementary figures 1-2**). Briefly, slides with initial human annotations were input into a U-Net convolutional neural network (CNN). The resulting algorithm generated probability masks and object segmentations, which were then reviewed by human experts to identify overlooked or wrong objects and modify incorrect annotations. This phase involved a thorough object-by-object verification of both human- and AI-annotated objects (**Figure 1B**). The generated algorithms were also applied to new, unannotated slides to increase the number of generated annotations and to take into account the variations in the histological context.

As a further (and final) step, in order to improve the accuracy and to speed-up the human-machine correction iterations, we developed a Dice score guided object correction. Annotations obtained after the three initial rounds of human-AI adjustments were used to train a 4th U-Net algorithm. Each annotated and/or AI-detected object was individualized, and a Dice score was calculated. Individual objects and associated Dice scores were uploaded to the integrated Calculator module of Visiopharm. This module allows at the same time to visualize the Dice scores of all individual objects and the corresponding objects’ outlines on the annotated slide (**Figure 1C**). Using this procedure, we focused the final corrections only on objects with suboptimal Dice scores (<0.75) to optimize the workload for the human corrector.

In addition to creating ground truth datasets, we also evaluated strategies to further reduce, in the future, the annotation burden for human experts during initial labeling of objects. We tested several options for the objects’ outlining simplification and evaluated the different conditions on AI-based automated segmentation (see **Supplementary data** and **supplementary Figure 3**).

### Algorithm training and testing for object detection and segmentation

After creation of the ground truth using the iterative AI guided corrections, two distinct algorithms were trained for the detection and segmentation of NPs and NFTs.

For the NPs algorithm, we divided our initial set of fifteen annotated slides in a training set containing twelve annotated slides, bearing a total of 4643 annotated and expert-verified NPs, and a test set with three slides which included a total of verified 1133 NPs (**Supplementary Figure 1, Supplementary table 2**). The detection/segmentation algorithm was generated in Visiopharm® using available U-Net CNN applied on WSI with 100x magnification and a 1024x1024 pixels field of view. In addition to this, a data augmentation step, including image rotation, flip, changes in intensity and hue, was performed to increase the robustness of the training. After object segmentation, post processing steps were applied including object median smoothing, filling holes and filtering of non-relevant small objects (<50µm²).

For the NFTs algorithm, the slide set was composed of nine annotated and expert-verified slides, which were subdivided in seven slides for training (containing 4050 objects) and two slides for testing (1093 objects) (**Supplementary Figure 2, Supplementary table 2**). The detection/segmentation algorithm was created using 200x magnification and a 1024x1024 pixels field of view window. Data augmentation steps, including image rotation, flip, changes in intensity and hue, were applied. After object segmentation, post processing steps were performed including object median smoothing, filling holes and filtering of irrelevant small objects (<10µm²).

The training was stopped when the loss functions reached the value of < 0,005 for both algorithms. A separate application was created for the calculation of precision (i.e., positive predictive value) and recall (i.e., sensitivity) as well as F1 (i.e., object detection) and Dice (i.e., object segmentation) scores:

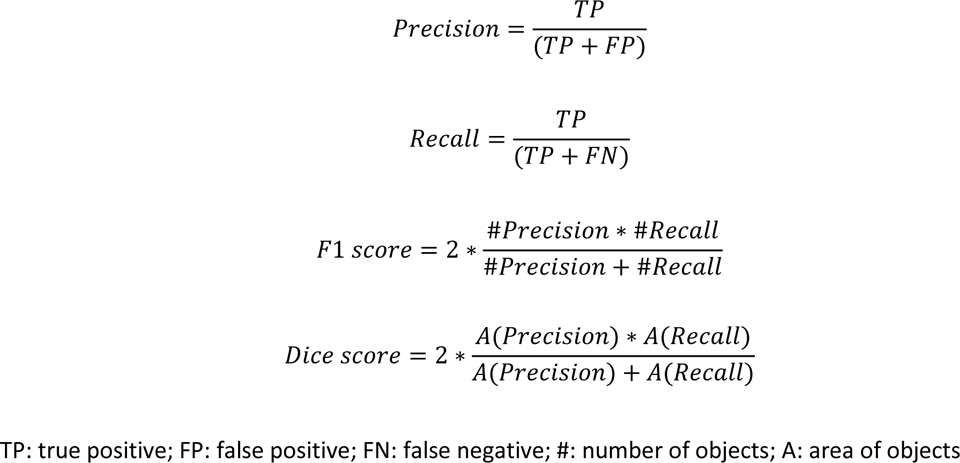

## Results

### Ground truth generation: 1) Enhancing annotation quality of neuritic plaques and neurofibrillary tangles through AI-driven iterative approach and object-by-object human validation

We used six slides with different densities of lesions to generate ground truth for NPs objects. The goal was mainly to evaluate the impact of a U-Net based model assistance on the discovery of true objects that were overlooked by human annotators. Two experts (LS, LI) manually drew 1069 annotations of NPs, agreeing on both the location and outline of the objects. This human-annotated set was used to train a first model, the output of which was then validated or corrected by human annotators (LS, LI, SB). The AI assistance and Human-in-the-loop allowed to identify objects initially overlooked by experts. Three subsequent rounds of AI training and human corrections were performed, using the initial set of slides, which yielded an increase of the number of expert-validated NPs by 43.9% after three correction rounds compared to the initially hand-annotated dataset (Database 1). In addition to the six initially annotated slides, nine new (non-annotated) slides were included in the annotation-correction procedure to constitute a 5013-objects expert validated ground truth dataset for NPs.

We applied the same procedure to NFTs annotations, starting with five slides presenting variable NFTs burden. This dataset included 1939 human-annotated and verified NFTs annotations. Training a first U-Net model and performing subsequent three rounds of AI-Human iterative validation/correction yield 2325 NFTs annotations, representing an improvement of 19,9% of segmented objects compared to the initial “human-only” annotated dataset. In addition to the five initially annotated slides, four new (non-annotated) slides were included in the annotation-correction procedure to constitute a 4927-objects expert validated ground truth dataset for NFTs.

Our iterative process demonstrated the importance of combining the strengths of AI and human experts to achieve a more accurate and comprehensive dataset of NPs and NFTs, providing a valuable resource for further research in the field of digital neuropathology.

### Ground truth generation: 2) Dice scores filtering allows fast and precise annotation dataset correction

As a supplementary and final step, we used individual object Dice scores (for NFTs or NPs) to guide and optimize annotation correction. We collected individual Dice scores for each annotated object and first corrected objects with the lowest Dice scores (0 value), corresponding to AI-segmented but not human-annotated objects. This included the false positive objects (i.e., objects that were recognized as neuritic plaques by AI but do not belong to this category according to the pathologist) as well as “overlooked” objects (i.e., objects correctly recognized as neuritic plaques by AI and that were missed by the pathologist). These “overlooked” objects were outlined and added to the object dataset, leading to an 18.8% (p<0.0001) reduction in null Dice score objects (**Figure 3**). Objects with low (0.05 - 0.45) and moderate (0.45 – 0.75) individual Dice scores represented sub-optimal AI recognition and were manually corrected by pathologists. However, applied corrections led to only minor improvement in NPs detections (+4.1% for low Dice score objects and −1.6% for moderate Dice score objects). In contrast, there was a significant improvement (+18.3%; p<0.0001) in the number of “high agreement objects” (Dice score > 0.75) after human correction (**Figure 3A**). To summarize, Dice score-based corrections largely potentiated the congruence of human-AI segmentations of NPs (decrease of null Dice score objects and increase of high Dice score objects).

**Figure 3:**
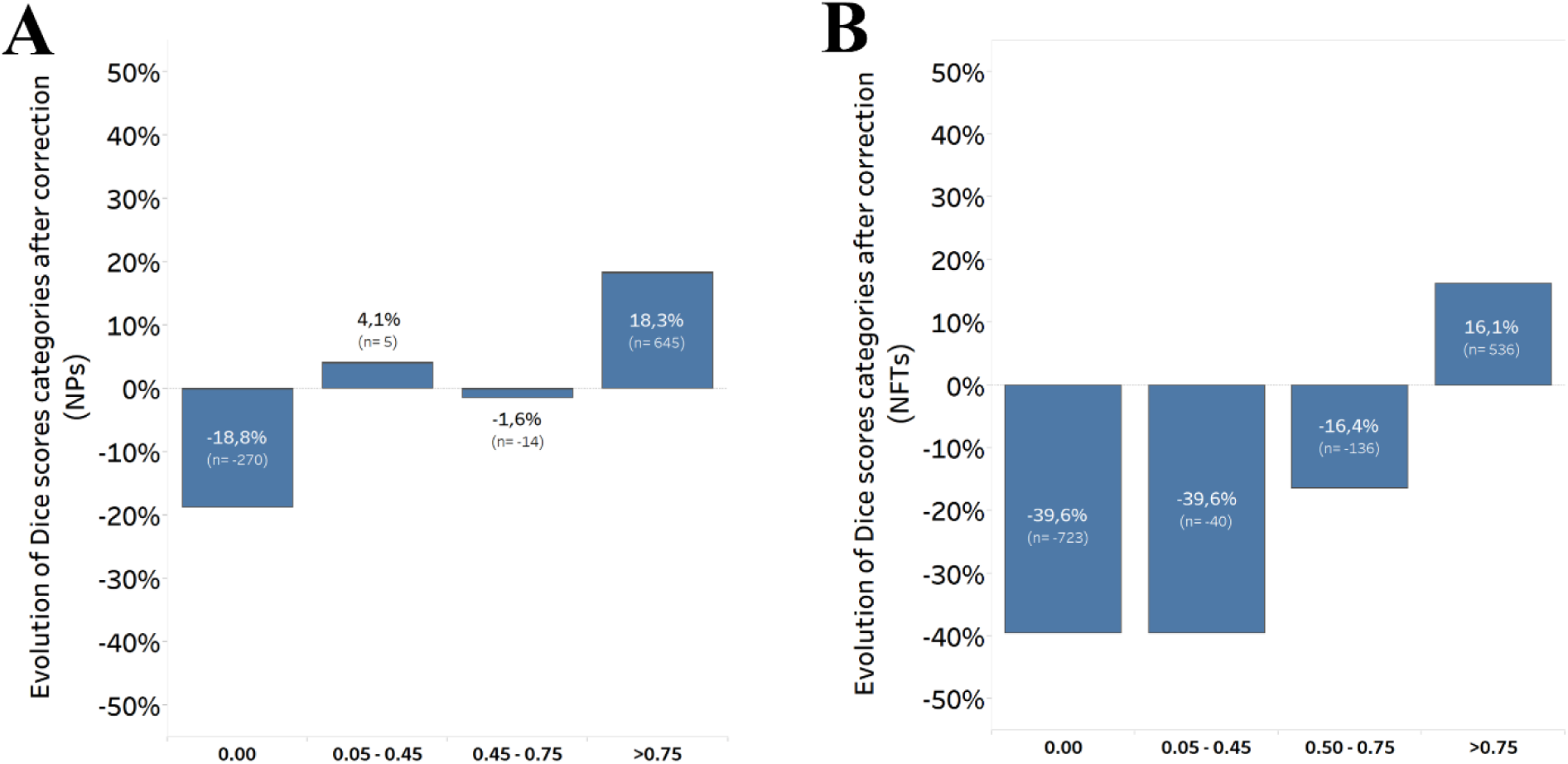
Number of objects with null, low, moderate, or high Dice scores following selected object correction based on initial Dice scores. The plotted values correspond to differences in counted objects in each Dice scores categories (post-correction minus pre-correction). **A:** Improvement of the segmentation accuracy after the Dice score guided annotation correction of NPs. Note a nearly 20% decrease in the number of objects with a “0” Dice score and at the same time a large improvement of “well recognized” objects (Dice scores >0.75). **B:** Improvement of the segmentation accuracy of NFTs after the Dice score correction. Note an important decrease of the Dice score “0” and low Dices score objects (0.05-0.45), both by nearly 40%. Objects with moderate Dice scores (0.5-0.75) were also reduced. Concomitantly, the precision of the segmentation of objects with highest Dice sores (>0.75) was improved by more than 15%.

Similar to NPs, selection by Dice score had a high impact on “0” Dice-scored NFTs, significantly reducing this category by 39.6% (p<0.0001). Additionally, the relative number of low and moderate Dice score NFTs was also reduced by 39.6% and 16.4%, respectively while the number of high Dice score NFTs significantly increased by 16.1% after correction (p<0.0001; **Figure 3B**). As for NPs, Dice score-based corrections applied to NFTs objects allowed to strongly maximize congruence of human-AI segmentation (decrease of null Dice score objects and increase of high Dice score objects).

In conclusion, by correcting objects with low and moderate Dice scores and adding “overlooked” objects to the dataset, we were able to substantially enhance the robustness of the NPs and NFTs databases. This approach decreased the time and effort required for expert correction, thereby increasing efficiency.

### Ground truth generation: 3) improved annotation strategy

In parallel with our main study, we evaluated strategies to reduce the annotation burden on pathologists through outline simplification. We compared the accuracy of U-Net based algorithm segmentation of neuritic plaques derived from human annotations with precise, rectangular, or larger outlines. Our results (see **Supplementary data**) demonstrate that simplified rectangular outlined objects yielded high accuracy and correctly segregated peripheral background and object limits with the same performance as highly-detailed and precise manual outlines. In contrast, algorithms trained with larger outlined objects were less accurate. These findings suggest that a simple bounding-box or simplified polygonal annotations can provide enough information for AI-segmentation of inner objects while saving time and effort for human annotators.

### Segmentation of neuritic plaques (NPs) using AI-improved object database

To segment neuritic plaques (NPs), we used our training set of twelve annotated whole-slide images (WSIs) containing 4643 expert-validated NPs (see part (2) of supplementary Figure 1). We implemented a U-Net detection and segmentation algorithm, which achieved a loss function of 0.003 after 95,000 iterations. The trained algorithm was then tested on three additional WSIs containing 1133 annotated objects.

The detection capacity of the algorithm was evaluated using the F1 score. NPs were detected with a mean F1 score of 0.86 (±0.04). The algorithm was evaluated on a test set of 1133 annotated NPs. It detected 1079 objects, of which 959 were true positives (TP) and 120 were false positives (FP). The recall and precision of the algorithm for NPs detection were 0.86 and 0.89, respectively, indicating a low rate of both false negative and false positive errors.

The segmentation accuracy was evaluated using the Dice score and achieved a value of 0.77. Of the total segmented area, 81% was correctly classified as belonging to the NPs category (precision: 0.81). Of the total annotated NPs area, 73% of segmented pixels were correctly classified as belonging to this category (recall: 0.73). These data are summarized in **Table 1**.

**Table 1:**
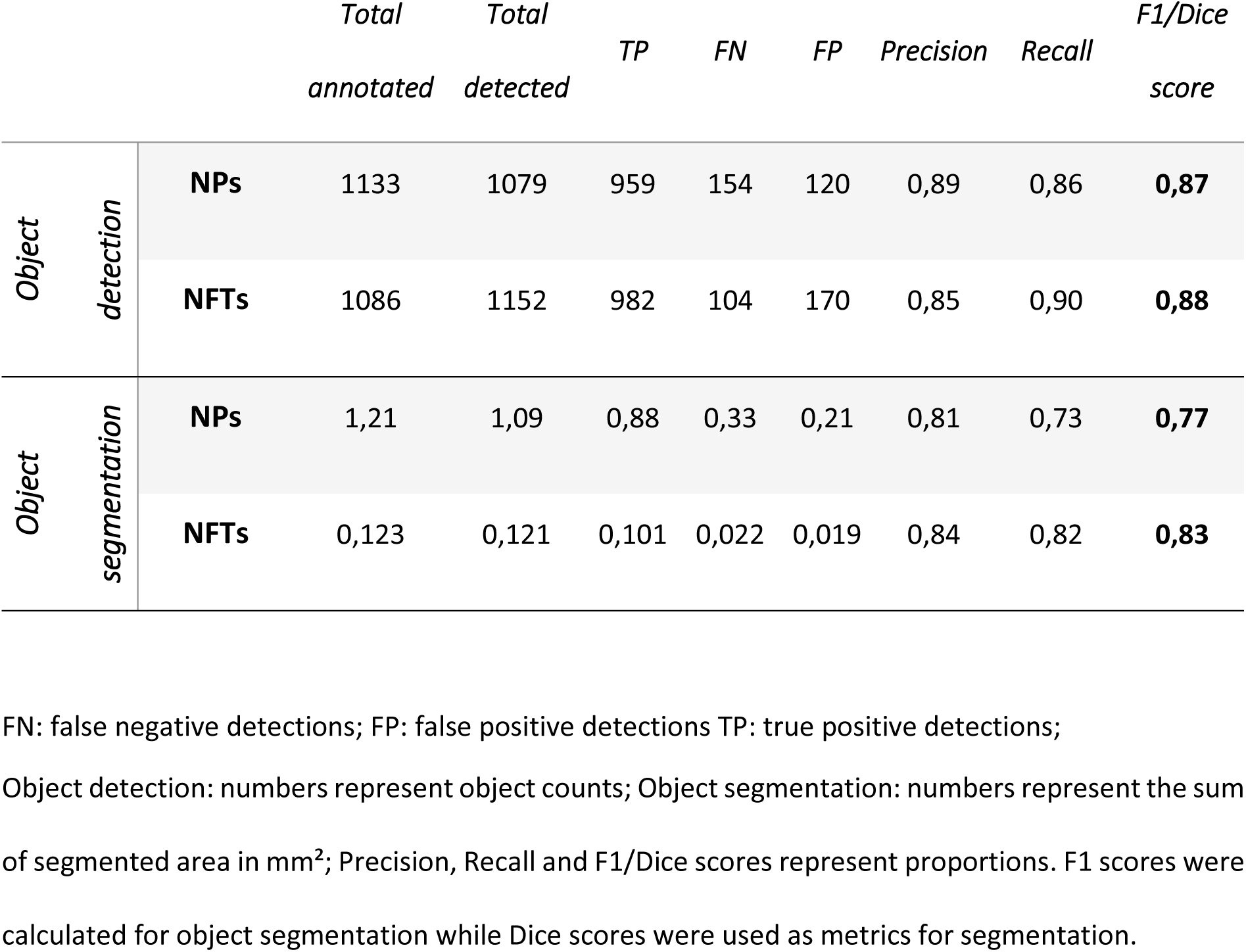
Detection and segmentation metrics for neuritic plaques and neurofibrillary tangles data sets.

In summary, our algorithm allowed for high-precision detection and segmentation of NPs, enabling calculation of number, load, and density as well as numerous morphology-associated measurements that could be derived from object segmentation.

### Segmentation of neurofibrillary tangles (NFTs) using AI-improved object database

For training the detection and segmentation algorithm using a U-Net based CNN model, 4050 objects were annotated on a set of seven slides (see part (2) of supplementary Figure 2). The model underwent more than 59000 iterations of training to achieve a final loss function of 0.004. The model was then tested on two supplementary annotated slides containing a total of 1086 NFTs. The algorithm detected 1152 objects, of which 982 were true positives (TP) and 170 were false positives (FP). The F1 score was 0.88 for NFTs detection. The precision and recall of the algorithm were 0.85 and 0.90, respectively, indicating a low rate of both false negative and false positive errors. The accuracy of segmentation was evaluated using the Dice score which resulted in a score of 0.83. A precision and recall for segmented area belonging to NFTs category were achieved at rates of 0.84 and 0.82 respectively.

In conclusion (**Table 1**), our algorithm demonstrated precise segmentation capabilities for NFTs within the evaluated slide set. This enabled accurate evaluation for both number and morphological properties of NFTs.

## Discussion

### Main objectives

Our work specifically aimed to develop and evaluate deep learning-based image models for NPs and NFTs segmentation using phospho-tau (AT8) immunostained brain tissue from the routine neuropathological diagnostic workflow. We demonstrated that U-Net-based artificial intelligence algorithms, trained from extensive and highly controlled datasets, allow detection and segmentation of NPs and NFTs with impressive precision despite high morphological heterogeneity of sampled objects.

### Building reliable datasets for training DL models

Variations in object morphology combined with technical issues promoting heterogeneity in histological preparations (e.g., tissue conservation, stain intensity, artifacts etc.) represent a major challenge for efficient generalization of AI-based segmentation protocols. These variations are exacerbated when comparing images across institutions that do not use identical protocols for tissue preparation and staining. Several initiatives have therefore been launched to create online databanks containing whole slide images (WSI) that can be used to train AI-based algorithms to exclude artifacts and minimize the impact of color variations [8, 38].

In our study, we first addressed the challenge of creating a robust and extensive annotation dataset for complex objects like NPs and NFTs. We derived training datasets from slides, obtained from four different European brain banks, and presenting large heterogeneity in terms of density of immunostained objects and overall staining levels.

The success of AI-based segmentation algorithms is directly dependent on the quality of ground truth annotations [18]. However, even overtrained, and experienced human annotators (e.g., pathologists) face challenges when using WSI due to the large number of required annotations and difficulty in explicitly annotating all objects of interest in a given area. This caveat becomes critical when performing annotation work in densely stained WSI comprising thousands of objects, as often seen in AT8 stained brain sections of patients with late-stage AD. We hypothesized that an AI-driven iterative procedure could increase the precision and speed-up human-made annotations of dense and heterogeneous objects.

We developed an AI-based “human in the loop” approach using a commercially available image analysis platform (Visiopharm®) to improve the quality of annotations of neuritic plaques and neurofibrillary tangles. This method involved an iterative object-by-object correction mode utilizing a U-Net based algorithm trained on a small number of human-annotated objects. The algorithm produced suggestions for relevant objects (NPs or NFTs), which were then verified by a human annotator and included in subsequent training/correction rounds. This procedure resulted in a 43.9% and 19.9% improvement in annotation quality for NPs and NFTs, respectively.

However, this approach remains time consuming for the human annotator. To address this issue, we developed a subsequent Dice score guided correction method that allowed experts to focus their efforts on unannotated or “overlooked” objects (Dice score 0) and on objects with low segmentation quality (Dice score <0.45) suggesting bad agreement between AI and pathologists. This refinement made the correction process faster and less error-prone, leading to further improvements for both NPs (−18.8% for Dice score 0 objects) and NFTs (−39.6% for Dice score 0 objects) datasets. The resulting final training datasets were validated by two pathologists (LS, SB) and deemed suitable for final training of AI-based segmentation protocols. During training datasets generation, we confirmed that even trained human annotators can overlook many objects during the initial annotation phase, which may lead to imprecise datasets and propagation of errors to subsequent AI-based segmentation algorithms. To solve this issue, we showed that AI-based algorithms can effectively identify overlooked objects and significantly improve annotation quality to provide refined training datasets and establish a reliable ground truth.

### Automated detection and segmentation of plaques and tangles

Using our validated and extensive annotation datasets we finally trained two algorithms to detect and segment NFTs and NPs. Performance of the generated models was evaluated as high (**Table 1**).

Zhang et al. [39] and Signaevsky et al. [34] previously developed deep learning-based models for detecting and quantifying NFTs in AD and other tauopathies. In our study, we achieved slightly higher precision in NFTs detection (NFTs F1 score: 0.88) compared to Signaevsky et al. (NFTs F1 score: 0.81). In addition, and contrarily to previous studies, we were able not only to detect tangles but also to segment them within defined grey matter regions, a step that will permit to perform ensuing morphological measurements of individual tangles (e.g., analysis of metrics such as areas, shapes, textures etc.).

Recently, Koga et al. [15] successfully detected and classified neuritic plaques, astrocytic plaques, and tufted astrocytes using 250x250 pixel static images of phospho-tau positive objects submitted to the Google AutoML platform for training. The model allowed the automated identification of different tauopathies associated to the distinct tau aggregates that were detected. A high precision classification of NPs versus other phospho-tau positive objects was obtained, underlining that NPs present indeed specific features that enable their detection. However, once again, the segmentation of NPs was not implemented in the model as opposed to our algorithm.

### Future developments and applications

#### Optimization of training datasets generation

Object segmentation remains a challenging task for both human annotators and computer vision procedures [28]. Experts (i.e., pathologists) are assumed to have a fair understanding of segmentation when performing annotation work. However, this assumption is often challenged by the large number of images to be annotated and the optical crowding of objects in WSI. Several studies aimed therefore at simplifying the annotation process through interactive segmentation methods [28]. In medical imaging, non-expert or expert users were for instance asked to draw simple polygons around hip joints [5], muscle and melanoma cells [7], and nuclei [9]. While these approaches require further improvement to be applied to WSI, they aimed to enhance reduce annotation time while preserving ground truth quality and subsequent model training.

Neuritic plaques with complex geometries can be particularly challenging to outline. In the present study, we evaluated the impact of outline simplification on the accuracy of a U-net based segmentation algorithm. Our results indicate that the algorithm was only weakly affected by variations in geometric precision and that even simple bounding-box annotations can provide sufficient information for AI-segmentation. Reducing the burden of objects outlining can dramatically improve annotation speed in future works.

#### Tailored open-source and explainable algorithms

From a technical point of view, one major disadvantage of commercial software is the user’s lack of traceability, interpretability, and explainability on the trained deep learning models [29]. Therefore we started testing in-house traceable tools for NFT and NPs segmentation and detection [13]. These tools, also derived from the dataset used in the present study, are based on CNNs and UNet architectures. To enhance the segmentation performance, we used attention mechanisms that suppress irrelevant regions in the input patches and highlight salient features of the NFTs and NPs. We evaluated our tools using cross-validation and cross-testing, which ensure the robustness and generalization of our results followed by the use of visual explanation methods, such as attention maps, to provide insights into the segmentation process of the deep learning models. Our traceable tools achieve high Dice scores (up to 73%) for NPs segmentation, after cross-validation and without any postprocessing[13]. Following these initial results, we plan 1) to implement the same approach for the segmentation of NFTs and 2) to evaluate other visual explainability algorithms, such as the SHAP [19] or GradCAM [33].

#### Biological applications

Our long-term goal is to stratify Alzheimer’s Disease patients based on fine and automated histopathological analysis using tools such as the ones implemented in the present study.

Deep learning models may be useful in segregating, on histological grounds, different subtypes of Alzheimer’s Disease (AD). Our next step will be to study cohorts of patients to determine if the morphological characteristics of neuritic plaques and neurofibrillary tangles vary between regions of interest (in the same individuals), if their geometry changes depending on the patient, if morphological heterogeneity allows stratification of patients and what morphological features are most significant and robust in distinguishing patients with different AD subtypes. Such analysis would be fruitful when applied to AD subtypes that, from standard neuropathological assessment, do not present obvious differences (e.g., classical vs rapidly progressive variants).

Having a pipeline to segment and differentiate tau aggregates will also participate, in the long run, to establish reliable diagnostic, prognostic, and associated treatment options.

#### Conclusions

In conclusion, our data indicate that artificial intelligence-based algorithms can be effectively applied to histological material for two main purposes. First, these algorithms can be used to identify overlooked objects in tissue sections and significantly improve the quality of annotations for creating labeled datasets (ground truth). Second, they allow to generate deep learning models capable of detecting and segmenting discrete brain lesions (such as plaques (NPs) and neurofibrillary tangles (NFTs)) with high precision in an unbiased and automated way.

## List of abbreviations

AD: Alzheimer’s disease
AI: Artificial intelligence
CNN: convolutional neuronal network
FN: false negative
FP: false positive
NFTs: Neurofibrillary tangles
NPs: Neuritic plaques
TP: true positive
WSI: whole slide image.

## Declarations

### Ethics approval and consent to participate

Informed consent was obtained from all subjects involved in the study.

### Consent for publication

All authors agreed to publish the present scientific work.

### Availability of data and material

Data that support the findings of this study are available from the corresponding author, upon request.

### Authors’ contributions

LS, BD, DR: drafting the study; LS, SB, LI: preparation and evaluation of annotations; GJ, AK: verification of AI-based segmentation algorithms.

## Acknowledgements

We are grateful to the patients and their families. We thank the Histomics core facility staff of the Paris Brain Institute for technical help for slide preparation and digitalization.

## Authors*’* information (optional)

## Supplementary data

### Evaluation of the impact of object outline precision on the accuracy of U-Net based segmentation

Complex objects, as neuritic plaques, are often difficult and fastidious to outline because of their complex geometries. These naturally imprecise limits can lead to a large inter-individual variation in annotation precision, related to differences in exact object outline. Thus, to obtain a robust annotation dataset several verification rounds and a strong consensus between pathologists are required. In order to know if we could simplify and speed-up the human annotation creation, we tested various protocols of simplification of object outline and evaluated the impact of these simplifications on the accuracy of U-Net based segmentation algorithm (**Figure S3**). A training set containing 12 slides with 4013 neuritic plaques was used for training. Obtained algorithm was tested on three different slides with 1000 objects (**Supplementary table 2**). The first condition “Normal outline” represented the most accurate (but very time consuming) manual outlines of neuritic plaques. In the second condition (“Rectangular outlines”), these annotations were replaced by smallest enclosing rectangles (i.e., bounding box around the object). The third condition consisted in “Bigger outlines”, in which normal outlines were grown by 10 pixels. In the fourth condition (“Bigger rectangular outlines”), normal outlines were replaced by the smallest enclosing rectangle that was grown by 10 pixels. Each set of objects was used for training of a specific segmentation algorithm. Obtained segmentations of each condition were compared to the “Normal outlined” objects used as ground truth and a general Dice score was calculated for each condition (with normalization to “Normal outline” Dice scores. The different scores were submitted to statistical analysis: One-way ANOVA followed by Dunnett’s multiple comparisons test was performed using GraphPad Prism (version 9.4.1 for Windows, GraphPad Software, San Diego, California USA).

Results indicate that algorithm trained with rectangular outlined objects presented a high segmentation accuracy illustrated by a relative Dice score of 0,96 with no difference to the “Normal outline” condition (p=0,168; **Figure S3B**). This algorithm, even with rectangular annotations as training data, produced non-rectangular segmentations. This shows that it could accurately separate the background and the object boundaries. Segmented objects strongly resembled normally outlined objects with only occasional imprecision of the outline. Algorithm trained with “Bigger outlined” objects was on the contrary less accurate (relative mean Dice score: 0,86) and the outlines were significantly less precise than those of the reference group (p=0,002). The last algorithm, trained with bigger rectangular outlined objects, presented the lowest accuracy (relative mean Dice score: 0,74). In this latter condition, segmented objects were significantly larger compared to the normally outlined objects (p<0,0001), but remained of similar shape.

In summary, the obtained results indicate that used U-Net based algorithms can provide high accuracy even when trained with highly simplified annotations (smallest rectangular bounding boxes). This opens future strategies to minimize human annotation burden to build training datasets.

### Supplementary Figure

**Supplementary Figure 1:**
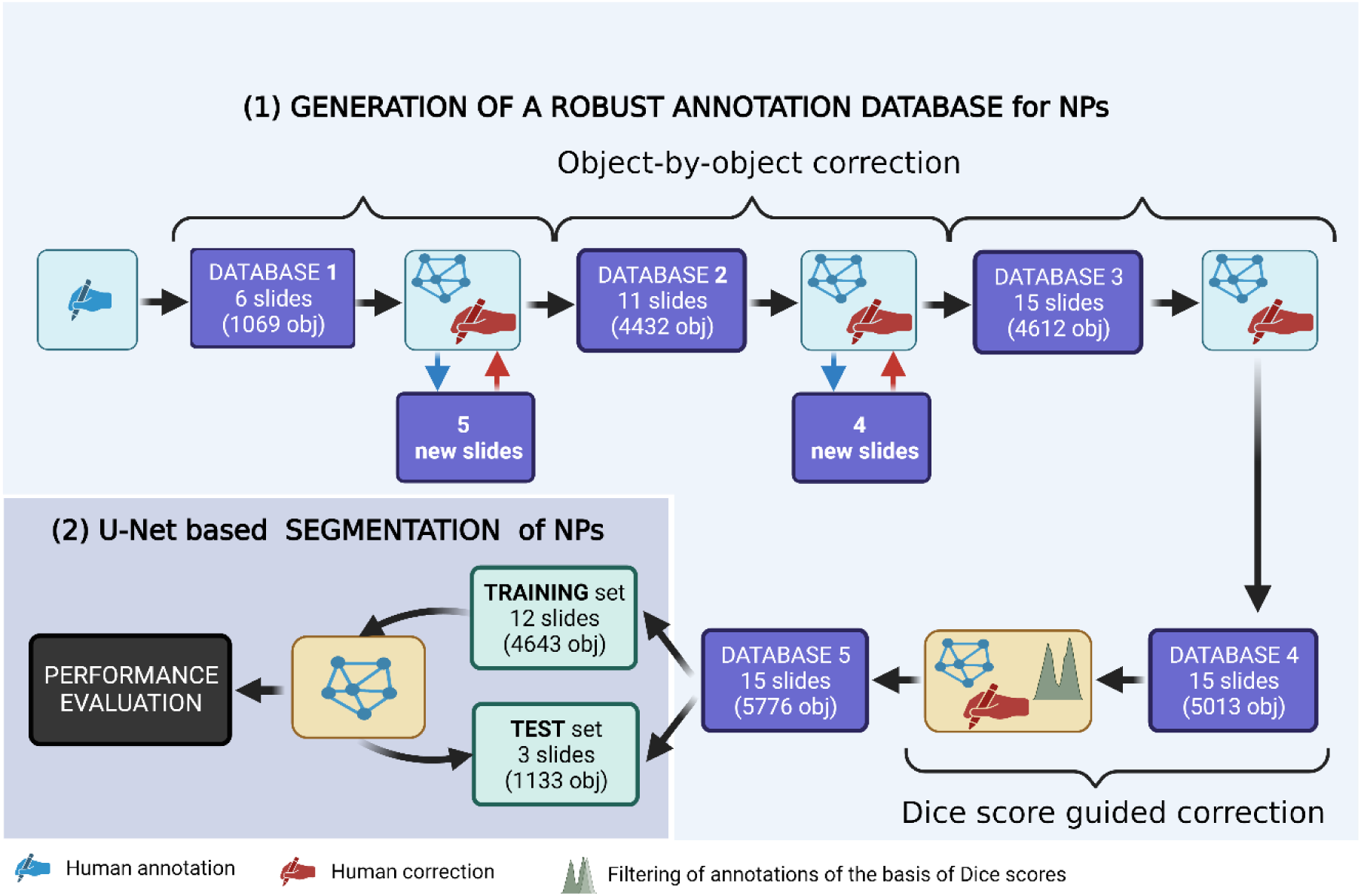
Workflow of annotation and correction of NPs. Presented workflow used object-by-object and Dice score correction methods (1) and training/test design for generating the DL model to detect and segment NPs (2).

**Supplementary Figure 2:**
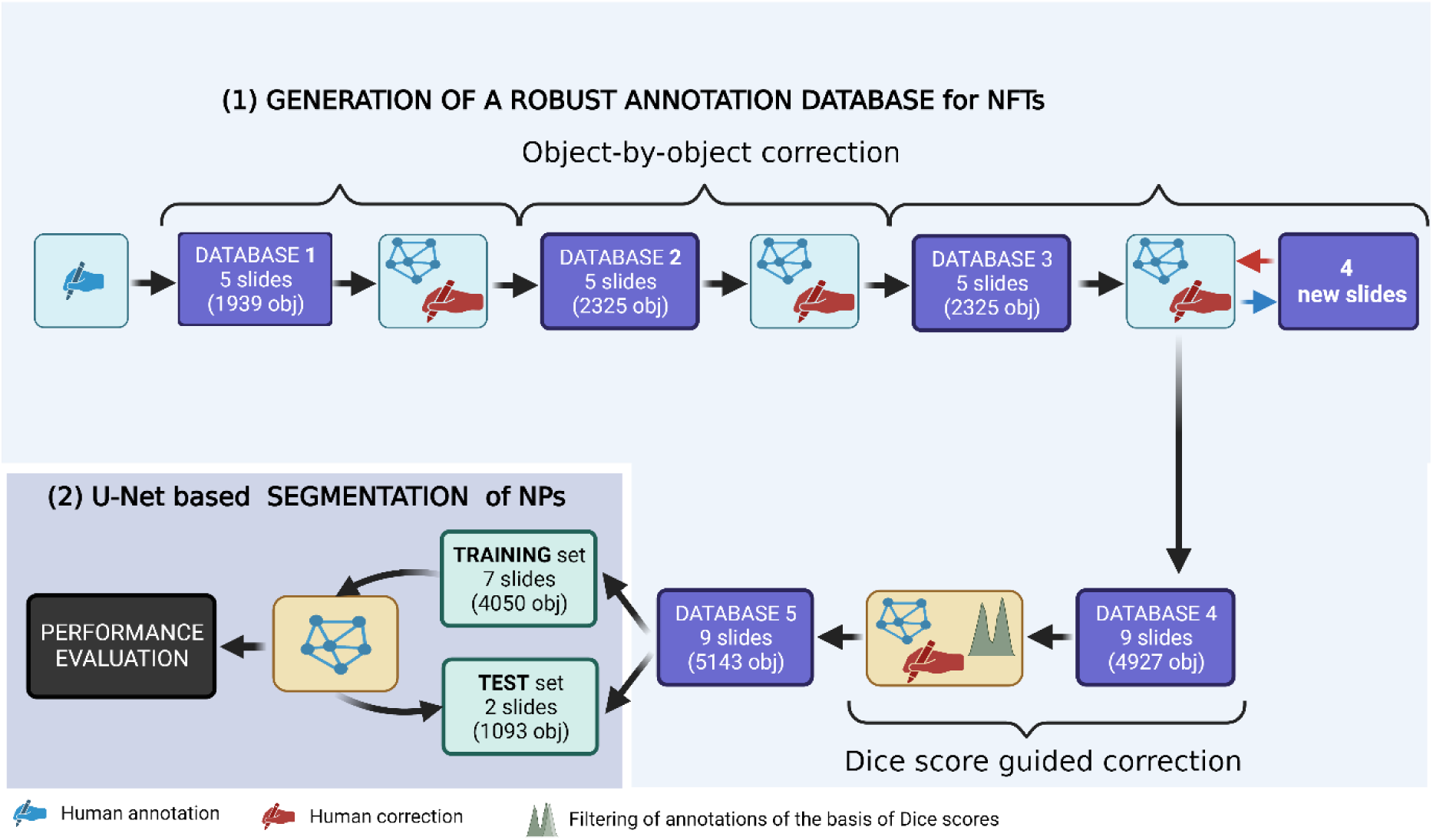
Workflow of annotation and correction of NFTs. Presented workflow used object-by-object and Dice score correction methods (1) and training/test design for generating the DL model to detect and segment NFTs (2)

**Figure S3:**
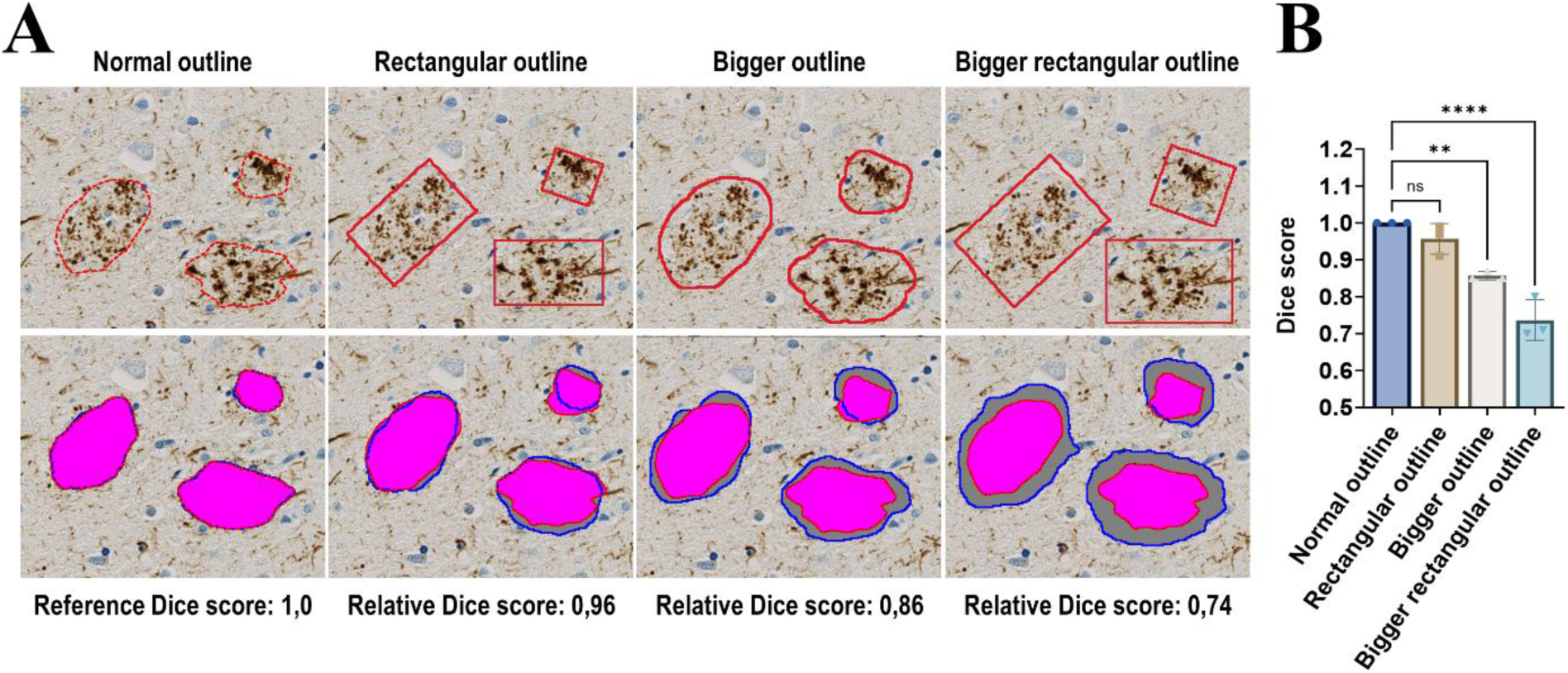
AI-object segmentation precision relies on human annotations’ outlines. **A:** Four modalities of object outlines (here NPs) were evaluated. First line illustrates the different types of object annotations. Second line corresponds to the AI-generated segmentation masks (grey surfaces with blue outlines) overlayed with ground truth annotations (pink surfaces with red outlines). Note the overall fidelity of segmented surfaces generated from rectangular outlines compared to “Bigger outline” and “Bigger rectangular outline” conditions for which AI-segmented areas were clearly overestimated. **B:** Quantification of the relative Dice scores in the different conditions. ns: non-significant, ** p < 0.001, **** p < 0.0001.

**Supplementary Table 1:**
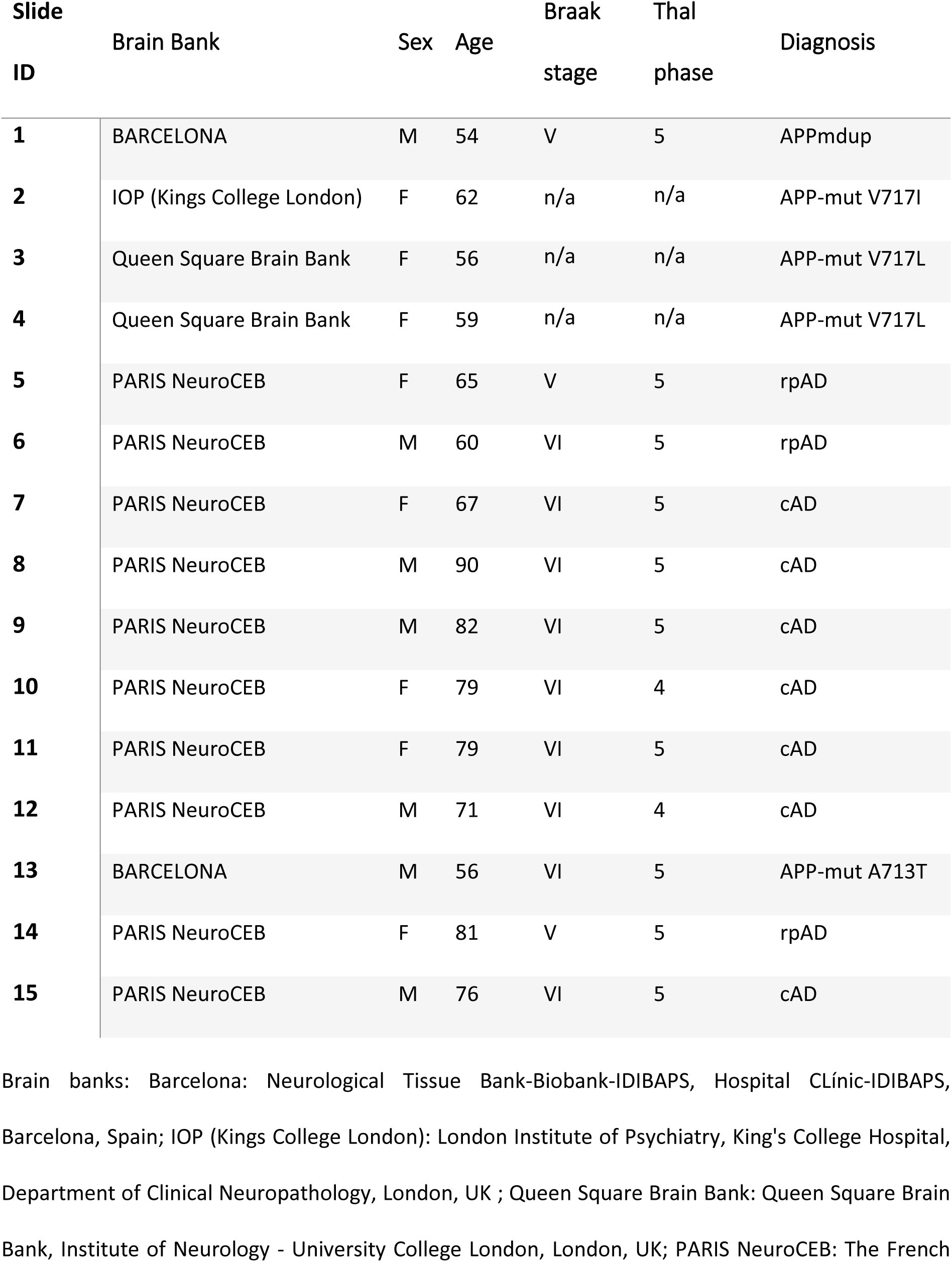
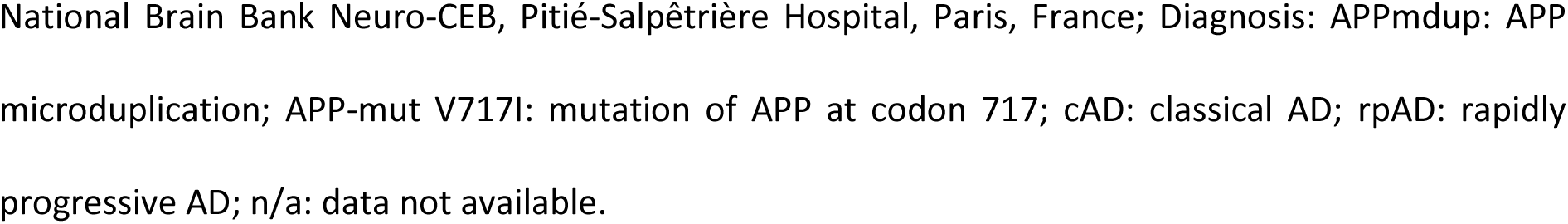
Clinical, genetic, and neuropathological characteristics of cases provided for the study.

| Slide ID | Brain Bank | Sex | Age | Braak stage | Thal phase | Diagnosis |
| --- | --- | --- | --- | --- | --- | --- |
| 1 | BARCELONA | M | 54 | V | 5 | APPmdup |
| 2 | IOP (Kings College London) | F | 62 | n/a | n/a | APP-mut V717I |
| 3 | Queen Square Brain Bank | F | 56 | n/a | n/a | APP-mut V717L |
| 4 | Queen Square Brain Bank | F | 59 | n/a | n/a | APP-mut V717L |
| 5 | PARIS NeuroCEB | F | 65 | V | 5 | rpAD |
| 6 | PARIS NeuroCEB | M | 60 | VI | 5 | rpAD |
| 7 | PARIS NeuroCEB | F | 67 | VI | 5 | cAD |
| 8 | PARIS NeuroCEB | M | 90 | VI | 5 | cAD |
| 9 | PARIS NeuroCEB | M | 82 | VI | 5 | cAD |
| 10 | PARIS NeuroCEB | F | 79 | VI | 4 | cAD |
| 11 | PARIS NeuroCEB | F | 79 | VI | 5 | cAD |
| 12 | PARIS NeuroCEB | M | 71 | VI | 4 | cAD |
| 13 | BARCELONA | M | 56 | VI | 5 | APP-mut A713T |
| 14 | PARIS NeuroCEB | F | 81 | V | 5 | rpAD |
| 15 | PARIS NeuroCEB | M | 76 | VI | 5 | cAD |

Brain banks: Barcelona: Neurological Tissue Bank-Biobank-IDIBAPS, Hospital Clínic-IDIBAPS, Barcelona, Spain; IOP (Kings College London): London Institute of Psychiatry, King's College Hospital, Department of Clinical Neuropathology, London, UK ; Queen Square Brain Bank: Queen Square Brain Bank, Institute of Neurology - University College London, London, UK; PARIS NeuroCEB: The French
National Brain Bank Neuro-CEB, Pitié-Salpêtrière Hospital, Paris, France; Diagnosis: APPmdup: APP microduplication; APP-mut V717I: mutation of APP at codon 717; cAD: classical AD; rpAD: rapidly progressive AD; n/a: data not available.

**Supplementary Table 2:** Slides used for building the initial object annotation and correction sets (A), for evaluation of object outline simplifications (B), as well as for training of final detection-segmentation algorithms (C).

| Slide ID | (A)<br>Initial annotations |  | (B)<br>Simplification of object outlines | (C)<br>Final database |  |
| --- | --- | --- | --- | --- | --- |
|  | NP | NFT | NP | NP | NFT |
| 1 |  | X | Training | Training | Training |
| 2 |  |  | Training | Training |  |
| 3 |  |  | Training | Training |  |
| 4 |  |  | Training | Training | Test |
| 5 |  | X | Training | Training | Training |
| 6 |  | X | Training | Training | Training |
| 7 | X | X | Training | Training | Training |
| 8 | X |  | Training | Training | Test |
| 9 | X |  | Training | Training | Training |
| 10 | X |  | Training | Training | Training |
| 11 | X |  | Training | Training |  |
| 12 |  |  | Training | Training |  |
| 13 |  |  | Test | Test |  |
| 14 |  |  | Test | Test |  |
| 15 | X | X | Test | Test | Training |

NP: neuritic plaques; NFT: neurofibrillary tangles

